# Modulation of predictive coding in auditory paradigms of varying complexity in children with Specific Language Impairment

**DOI:** 10.1101/2025.07.02.662722

**Authors:** Francisco J. Ruiz-Martínez, Elena I. Rodríguez Martínez, Brenda Angulo Ruiz, Ana Gómez Treviño, Raquel Muñoz Pradas, Carlos M. Gómez

## Abstract

Specific Language Impairment (SLI) is a persistent difficulty in the acquisition and use of expressive and/or receptive language, which negatively impacts academic and social development. The present study evaluated the validity of the statistical learning model proposed to explain language difficulties in children with SLI. To this end, two auditory paradigms of varying complexity, framed within predictive coding theory, were passively presented to children diagnosed with SLI and without neurological impairments. The paradigms consisted of stimulus sequences with decreasing or increasing frequencies, interspersed with the sporadic occurrence of unexpected tone endings. The psychophysiological response was recorded using EEG, focusing on the P1, Mismatch Negativity (MMN), Post-imperative Negative Variation PINV, and Contingent Negative Variation (CNV) components. Results showed an absent MMN and a higher P1 response to deviant tones in children with SLI, suggesting an impaired development of frontal MMN generators, possibly compensated by the primary auditory cortex. SLI participants also showed increased PINV and CNV responses during the most complex paradigm, which could imply greater cognitive effort and resource allocation to the reassessment of stimulus patterns in this group. Finally, the incomplete maturation of frontal areas in children of this age range (3 to 11 years old) was proposed to explain the increased activity observed in both groups for the N1/MMN elicited in the simplest paradigm. These findings support statistical learning as a valid model for understanding the possible neural basis of SLI, identifying this type of predictive EEG design as a potential early detection protocol.

## INTRODUCTION

The predictive coding is a theory of established scientific interest which proposes that our brain is continuously extracting patterns from the environment with the aim of constructing generative models based on the prior experience. The objective of this cognitive function would be anticipate future events minimizing the so-called prediction error, in order to constantly guide our attention and behavior (Friston, 2005; Winkler & Czigler, 2012; Heilbron & Chait, 2018).

Under this theory, the prediction error would be the neural mechanism involved in detecting the deviance between a prediction, determined by the previously perceived stimulation, and a new sensory input. This disparity would generate a psychophysiological response, with an intensity proportional to the deviation, aimed at updating the predictive model to enhance prediction accuracy (Friston & Kiebel, 2009; Friston, 2010; Hohwy, 2012; Brodski et al., 2015). In line with this, a broadly supported hypothesis proposes a Bayesian probabilistic model to explain the way in which the brain generate these predictions and update them to reduce the prediction error (Knill & Pouget, 2004; Doya et al., 2007; Gómez & Flores, 2011; Friston et al., 2017; Gómez et al., 2019).

The psychophysiological functioning of predictive coding and its potential impairment in clinical disorders has been broadly studied through EEG (González-Gadea et al., 2015; Fogelson & Diaz-Brage, 2021; Qela et al., 2025). For this purpose, neurobiological markers sensitive to psychophysiological impairments in clinical populations, such as the auditory P1 (Schall et al., 1997; Ruiz-Martínez et al., 2020; Sharma et al., 2005), the auditory mismatch negativity (MMN) (Näätänen et al., 2014; Erickson et al., 2016; Tada et al., 2019), the contingent negative variation (CNV) (Osborne et al., 2020; Lahoz et al., 2023; Prata et al., 2023), and the post-imperative negative variation (PINV) (Klein et al., 1996; Diener et al., 2009; Werner et al., 2011) have been frequently analyzed.

The auditory P1 is an early positive component elicited between 50 and 80 ms post-stimulus in the primary and secondary auditory cortices. Additionally, it may also exhibit frontal activation during tasks with higher attentional or cognitive demands, as a consequence of the involvement of thalamocortical projections. This potential is considered a reflection of initial and automatic sound detection, although it has been proposed to be modulated by active attention (Picton et al., 1974; Celesia, 1976; Alho et al., 1994; Herrmann & Knight, 2001; Ponton & Eggermont, 2001; Martin et al., 2008; Giuliano et al., 2014). The auditory P1 is not commonly evaluated in predictive paradigms due to its early elicitation, which generally precedes cognitive processes such as the expectations generated by predictive coding. However, in some studies, the P1 and even earlier positive components showed to be sensitive to stimulus omission or deviations in temporal stimulus onset (Bendixen et al., 2009; Schwartze et al., 2013; Lee et al., 2024).

The MMN is the most important neural marker of preattentive auditory discrimination. According to predictive coding theory, it reflects the prediction error generated by the violation of the expectation formed by a repetitive stimulus pattern (standard), due to the occurrence of an unexpected stimulus that changes the pattern (deviant) (Winkler, 2007; Garrido et al., 2009; Wacongne et al., 2011). This component consists of a negativity, usually elicited between 100 and 250 ms, with a predominantly frontocentral topography. It can be generated passively (without paying attention to the stimulation) and modulated by active attention (Näätänen & Alho, 1995; Sussman et al., 2003; Näätänen et al., 2012). Although it is widely associated with prediction error, the discussion about whether this component is a N1 modulation caused by fresh afferent activity of cortical neurons undergoing, undergoing an adaptation or habituation process, remains ongoing (May & Tiitinen, 2009; O’Reilly & O’Reilly, 2021).

The CNV is a slow negative brain potential commonly elicited by stimulus anticipatory paradigms, as it is fundamentally generated by the expectancy that a warning signal produces regarding the occurrence of a target stimulus (Walter et al., 1964; Brunia et al., 2011; Gómez et al., 2019; Kóbor et al., 2021). It is considered a reflection of cortical arousal and attention generated by response anticipation, mainly in motor tasks (Kropp et al., 2001; Kononowicz & Penney, 2016; Fattapposta et al., 2024), although it has also been suggested in passive studies to reflect sensory expectations (Mento, 2013; Ruiz-Martínez et al., 2022; Muñoz-Caracuel et al., 2024).

In some experimental settings, the CNV recovery to the baseline after the target onset is sustained over time, forming the so-called PINV component (Delaunoy et al., 1978; Kathmann et al., 1990). This component, usually associated with paradigms designed to induce in the subject a perceived lack of control over aversive stimuli, a sensation of uncertainty regarding the correctness of a response, or the reassessment of ambiguous contingencies (Rockstroh et al., 1979; Kathmann et al., 1990; Klein et al., 1996), has recently been elicited in adults by our research group using the same experimental protocol, suggesting a PINV role in predictive coding (Ruiz-Martínez et al., 2022). Both the CNV and PINV topographies are predominantly located in the frontal region, although the CNV is also generated in central and parietal areas (Cui et al., 2000; Gómez et al., 2001; Bender et al., 2006; Diener et al., 2009).

These components will be analyzed in the present research in a sample of children with Specific Language Impairment (SLI) and without neurological pathologies (ND: normodevelopment) through a predictive coding approach, with the objective of studying the potential alterations of the statistical learning in language.

As described in the DSM-V (American Psychiatric Association [APA], 2013), the SLI would be included in a more general category called Developmental Language Disorders, consisting of a persistent difficulty in the acquisition and use of expressive and/or receptive language. To be considered SLI, these language limitations must occur in the early stage of development within proper social environments and without sensory, emotional, physical, or cognitive deficits (Villegas, 2002; APA, 2013). As a consequence of these language challenges, individuals with SLI often present delayed or impaired reading and/or academic development (Young et al., 2002; Pennington & Bishop, 2009). All of which can negatively affect their adolescence in multiple aspects (social, emotional, behavioural, etc.) (Wadman et al., 2011a, 2011b, 2011c), and face a reduced sense of well-being and life satisfaction in adulthood (Arkkila et al., 2008).

The genetic involvement in SLI has been amply researched (Bishop, 2002, 2009, 2013; Li & Bartlett, 2012), and many theoretical models have been suggested to explore its underlying mechanisms. Among the most relevant hypotheses is the temporal processing hypothesis of language, which proposes a desynchronization between neural activity and speech rhythms present in prosodic and phonological expression (Goswami., 2019). Another significant theory is the speed-of-processing limitation hypothesis, which puts forward a psychophysiological difficulty in distinguishing phonemes with short temporal transitions due to slower auditory processing (Tallal, 2004). Finally, the statistical learning model also stands out, suggesting a reduced ability to extract implicit language patterns in individuals with this disorder (Lammertink et al., 2017). This last theory is the approach followed in the present study and could underlie the SLI limitations in acquiring grammatical structures.

More specifically, the statistical learning model proposes that language is structured by phonological and syntactic dependencies that establish hierarchical links between words, where certain elements (dependents) are governed by others (heads), defining the distribution of grammatical and semantic roles across sentences. This model would suggest that SLI could present a deficit in the statistical processing necessary for establishing dependencies in adjacent and, primarily, non-adjacent terms in the sentence, through the sequential and probabilistic pattern extraction (Saffran et al., 1996, 2002; Gibson, 1998; Hsu et al., 2014).

Several psychophysiological studies have analyzed SLI using event-related potentials (ERPs) (Bishop & McArthur, 2005), suggesting that speech processing deficits and differences in the spatial distribution of attentional auditory resources are associated with a delayed P1 component (Shafer et al., 2007; An et al., 2022). A relationship has also been found between a reduced or absent MMN and attentional deficits in auditory stimulus discrimination (Shafer et al., 2005; Datta et al., 2010; Kujala et al., 2017). A reduction in the Late Positive Component (LPC), which is involved in post-lexical reanalysis and integration, has also been reported (Haebig et al., 2018). A delayed or absent N400, commonly interpreted as a semantic index of prediction error, has likewise been observed in individuals with SLI (Shafer et al., 2005; Sabisch et al., 2006; Haebig et al., 2018). Similarly, the absence or reduction of the Early Left-Anterior Negativity (ELAN) and the P600 may be related to deficits associated with SLI in the early detection of syntactic violations and in reanalysis or repair processes, respectively (Fonteneau & van der Lely, 2008; Royle & Courteau, 2014; Purdy et al., 2014; Haebig et al., 2017).

N400 and P600 components have been widely studied within the framework of Bayesian processing to evaluate the statistical learning model of language development, often using sentences with unexpected or incongruent elements (e.g., the cool water burned him) (Kutas & Hillyard, 1984; Herten et al., 2005; León-Cabrera et al., 2017; Wang et al., 2023; Michaelov et al., 2023). However, to our knowledge, no ERP study to date has investigated the SLI within a purely predictive paradigm. This limitation justified the present study, along with the need for an early diagnostic protocol to identify the SLI before language emergency, enabling timely interventions to reduce cognitive and social deficits. To this end, the P1, MMN, CNV, and PINV components were evaluated to detect potential alterations in the SLI through the presentation of two passive auditory paradigms of different complexity in children with this disorder and ND.

The following three hypotheses were proposed to explain how a potential deficit in statistical learning might influence the psychophysiological responses of SLI participants compared to the control group:

1. ND children were expected to show a higher amplitude for the deviant than for standard trials, primarily at the MMN latency (where this effect is most commonly observed), and secondarily also at the PINV. Conversely, the MMN amplitude for deviants would be attenuated in SLI participants.
2. A greater response to the first-order condition (high repetition of the same tones in the auditory sequences), compared to the second-order condition (low repetition of tones sequences), was expected for the P1 and MMN components in both groups. The direction of this effect would indicate the degree of maturation of predictive mechanisms and consequently the prediction error. Specifically, a higher response to the second-order condition would suggest a more adult-like maturation pattern, reflecting the recruitment of additional neural resources to extract more complex patterns. Conversely, a higher amplitude in the first-order condition would suggest incomplete development of these brain areas.
3. A higher amplitude was expected for the second-order condition in both the CNV and PINV, primarily in the SLI group. The increase in CNV amplitude would reflect enhanced preparatory process associated with greater cognitive effort, whereas the increase in PINV amplitude would reflect a heightened allocation of resources for stimulus pattern reassessment.

If confirmed, these hypotheses would support the early use of this passive EEG paradigm for diagnosing SLI in children, thanks to its non-invasive and child-friendly nature. This is especially relevant given the high prevalence of SLI, which affects between 1.4% and 16.2% of school-aged children worldwide (Tomblin et al., 1997; Villegas, 2022).

## MATERIALS AND METHODS

### Participants

A total of 86 subjects divided in a normodevelopmental (ND) and a clinical group diagnosed with Specific Language Impairment (SLI) were recorded in two experimental conditions of different complexity. The ND group was composed by 51 participants aged 3 to 11 years (19 males and 32 females, mean age in days= 2723.94 ± 625.03 SD, range 1739 - 4023 days old) and the clinical group by 35 participants aged 3 to 10 years (25 males and 10 females, mean age in days= 2484.49 ± 589.15 SD, range 1271 - 3916 days old). Due to excessive EEG artifacts in the recording of the complex paradigm, one subject from the clinical group was excluded from the analysis. Thus, the analyzed group in this paradigm was composed by 34 subjects aged 4 to 10 years (25 males and 9 females, mean age in days= 2520.18 ± 558.28 SD, range 1686 - 3916 days old).

The clinical group was recruited from the Unidad de Desarrollo Infantil y Atención Temprana (UDIATE: https://hospitalveugenia.com/udiate-atencion-temprana-desarrollo-infantil/) attached to the Hospital Victoria Eugenia, operating within the Spanish Red Cross, which is dedicated to the diagnosis and intervention of neurodevelopmental disorders in children. Inclusion in the study required a clinical report confirming language deficits, endorsed by language specialists from the clinical center. This confirmation was based on standardized assessments such as the Navarre Oral Language Test – Revised (PLON-R; (Aguinaga Ayerra, 2005), the Illinois Test of Psycholinguistic Abilities (ITPA; (Kirk & McCarthy, 1961)), the Clinical Evaluation of Language Fundamentals – Fifth Edition (CELF-5; (Wiig et al., 2013), the Kaufman Brief Intelligence Test (KBIT; (Kaufman & Kaufman, 2004), the Peabody Picture Vocabulary Test – Fifth Edition (PPVT-5; (Dunn, 1959), or a structured clinical interview conducted by language therapists in accordance with DSM-V or ICD-10 criteria. Before enrollment, otorhinolaryngologists confirmed normal auditory function using audiometric tests and, when needed, brainstem auditory evoked potentials, ruling out hearing impairment as a confounding factor.

ND participants were recruited from various schools in Seville, Spain. Parental reports confirmed the absence of learning difficulties, neurological disorders, or developmental delays in these children.

Non-verbal cognitive abilities were assessed using the Kaufman Brief Intelligence Test (KBIT) for 80 participants (due to complications during the evaluation, some participants were unable to perform the test). The raw KBIT scores met the criteria for normal distribution and equal variances. Statistical comparison of the raw scores (RS) did not reveal significant differences in cognitive performance between the ND group (M =23.79, SD = ±6.03) and the clinical group (M =22.41, SD = ±6.94), as indicated by a t-test (t(78) = 0.95, p = 0.35, d = 0.22).

Experiments were conducted with informed and written consent of each participant (and/or parents/carers), following the Helsinki Protocol (code: 0818-N-21). The study was approved by the Biomedical ethical committee from the Junta de Andalucía.

### Experimental Procedure

Auditory stimuli were delivered via two Dell A215 speakers positioned bilaterally to the participant’s head, at an intensity of 65 dB, measured using a Velleman DVM1326 sound level meter. Each auditory stimulus consisted of a 200-ms sinusoidal tone with 20-ms rise and fall times. Tones were generated using an online tone generator (https://www.wavtones.com/functiongenerator.php, last accessed April 15, 2019) and subsequently edited in Audacity 2.3.2.

The experimental paradigm was programmed in E-Prime 2.0 and included two conditions differing in sequence complexity: first-order and second-order. Each trial comprised a sequence of four tones (S1–S4; see Fig. 1). To control for neurologic activation effects in the auditory cortex, sequences with increasing and decreasing frequencies were counterbalanced across participants.

**Figure 1.**
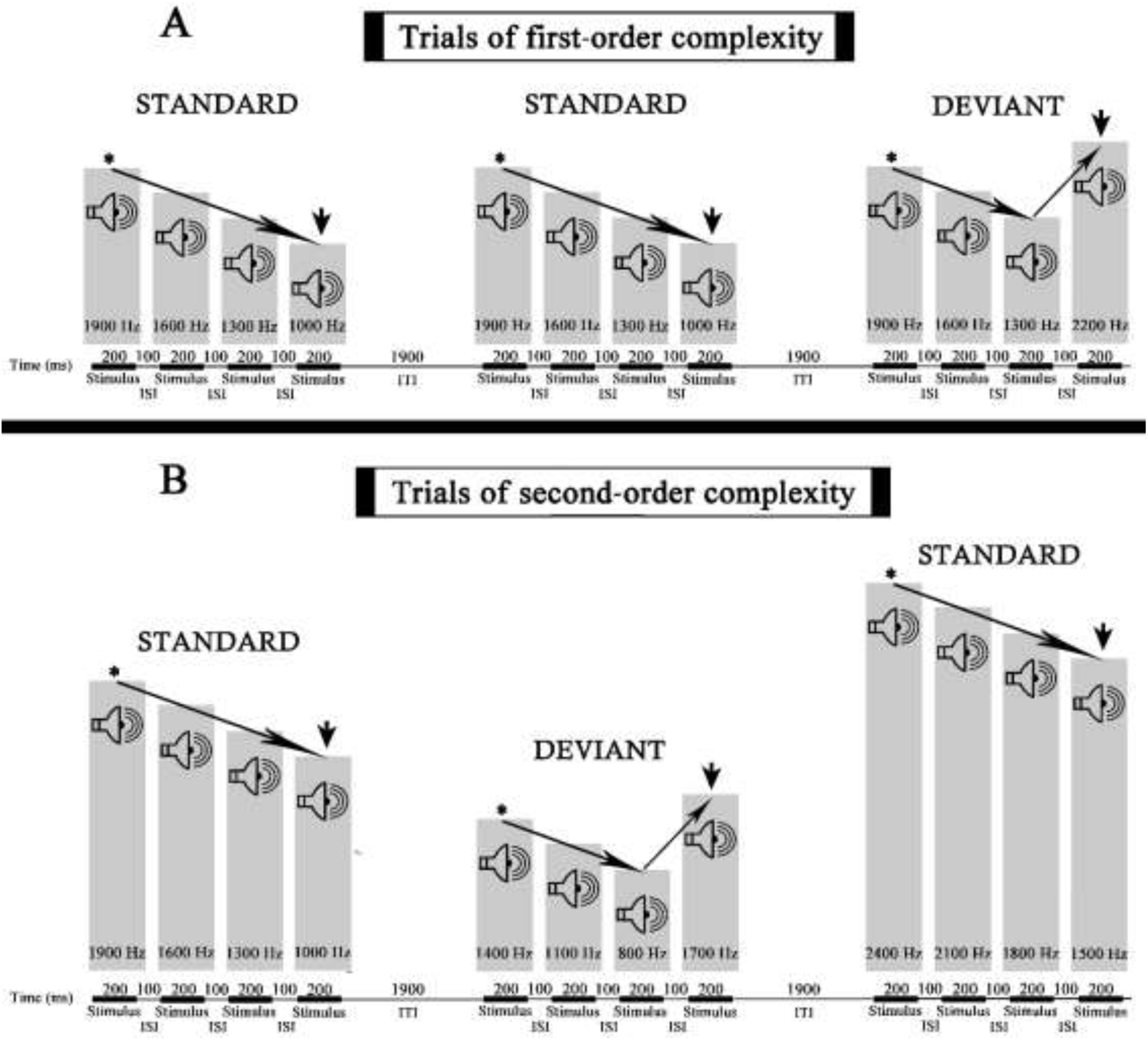
Experimental paradigms. A: the timeline below the bar graph shows the duration of the stimuli, the interstimulus intervals (ISIs), and the intertrial intervals (ITIs), the bar graphs present the order and frequencies of the stimuli displayed in the standard and deviant descending sequences for the first-order complexity. The fourth stimulus (S4), marked by arrows, defined the onset for time-locking P1 and late N1 epochs. Asterisks indicate that the baseline for obtaining the CNV and PINV components was located just before the 1st stimulus (S1).

Table 1 summarizes the frequencies used in both complexity conditions for ascending and descending sequences. In the first-order condition, each trial type (standard: S and deviant: D; ascending and descending) was defined by a single, fixed stimulus sequence, allowing participants to form stable representations of the physical characteristics of the tones.

**Table 1.**
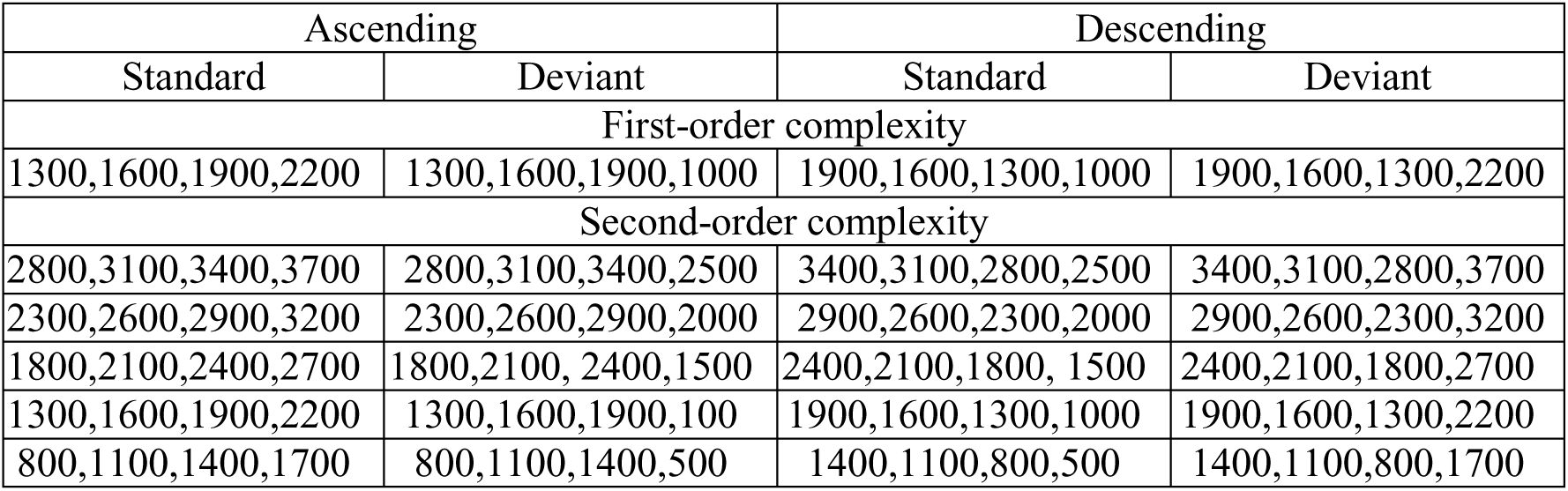
Stimulus frequencies by condition. The table displays the frequencies (in Hz) of each stimulus sequence presented in the ascending and descending sound designs, across both complexity conditions (first- and second-order) and trial types (standard and deviant).

In contrast, the second-order condition involved five distinct stimulus sequences per trial type, with a consistent frequency interval of 300 Hz between tones. Through this second-order condition, we examined if participants could detect the underlying frequency pattern while reducing the possibility from memorizing individual tone characteristics.

All trials had a total duration of 3 seconds, comprising four 200-ms tones, three 100-ms interstimulus intervals, and a 1,900-ms intertrial interval (Fig. 1). Each participant completed 604 trials in total: 241 S and 61 D trials per complexity condition, maintaining a 4:1 S-to-D ratio. The total experiment duration per participant was 30 minutes and 20 seconds.

The experimental blocks (first- and second-order) were presented in counterbalanced order across participants., trials were arranged considering the previous trial (SS, SD, DS, DD) with respective proportions of 59.1%, 15.7%, 15.7%, and 9.5% to ensure an adequate number of DD trials. The sequence of dyads was pseudorandomized such that at least one SS dyad followed any dyad containing a D trial, thus preserving the integrity of the S effect throughout the session.

### EEG recording

Signals were acquired from 20 scalp electrodes arranged according to the international 10–20 system (Fp1, Fp2, F7, F3, Fz, F4, F8, T3, C3, Cz, C4, T4, T5, P3, Pz, P4, T6, O1, O2), including a ground electrode. Given the potential sensory sensitivities in our participant population, we opted not to use ocular or mastoid electrodes. This decision facilitates replication across similar clinical groups and improvement of the quality of recordings by increased comfort of younger participants. A common reference was applied, and activity from electrodes Fp1, Fp2, F7, and F8 was used for detection and removal of blink and eye-movement artifacts.

Participants were seated comfortably between the loudspeakers and instructed to remain relaxed while viewing a silent movie displayed on a laptop screen throughout the recording session.

Data acquisition was performed in direct current mode at a sampling rate of 1024 Hz, using a commercial analog-to-digital acquisition and analysis system (ANT, The Netherlands) with an amplification gain of 20,000. Electrode impedance was kept below 10 kΩ, and data were filtered offline.

### Data analysis

EEG recordings were analyzed using EEGLAB version 14.1.1 and Matlab 2023b scripts that were developed in house. A high-pass filter of 0.05 Hz and a low-pass filter of 45 Hz were conducted on the EEG signal, in addition to an artifact subspace reconstruction (ASR) algorithm, implemented via the EEGLAB function “clean_rawdata”, which was employed to correct data segments whose S deviations exceeded 20 times the calibration data (Mullen et al., 2015).

We performed Independent Component Analysis (ICA) to identify and remove artifacts including those from electronic sources, muscle activity, cardiac signals, blinks, and eye movements in the epoched data. These components were removed and the EEG signal was reconstructed.

The mean of accepted components for the ND group was 12.63 ± 1.41 SD, range 9 – 16 (for the first-order complexity) and 12.59 ± 1.15 SD, range 11 – 16 (for the second-order complexity), whereas for the clinical group was 12.49 ± 1.65 SD, range 10 – 16, and 12.59 ± 1.89 SD, range 9 – 17, respectively.

Epochs were segmented using ERPLAB into different time windows, depending on the ERP component selected for analysis. This resulted in different averages for each participant, trial type (S or D), and complexity condition (first- or second-order complexity). Grand-average ERP images (Fig. 2 and 3) were generated to identify the ERP components, and to determine their respective latencies without introducing any condition or group bias. These figures present the activity recorded at each analyzed electrode across both participant groups and experimental conditions, in two different time windows.

**Figure 2.**
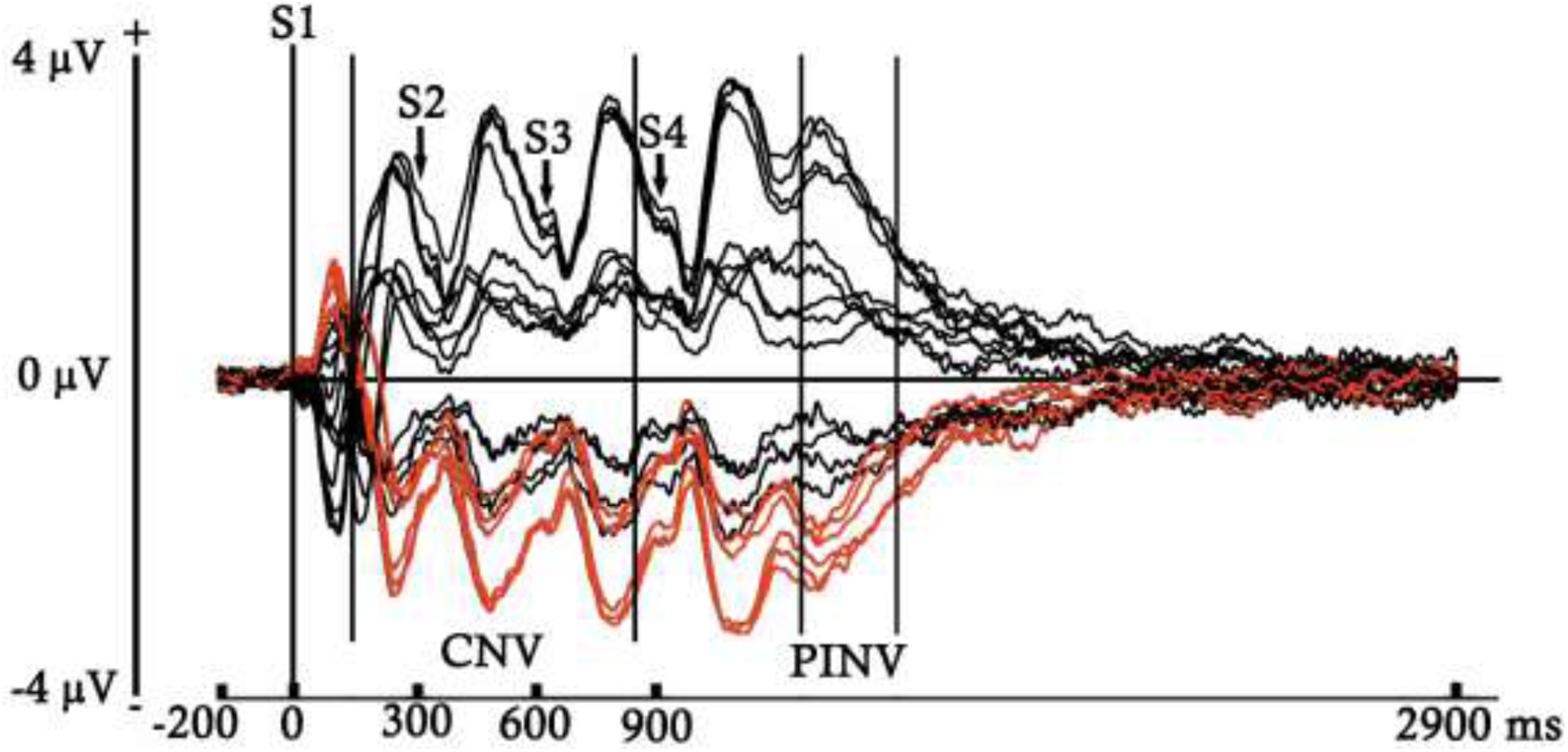
ERP averages obtained for both trial types (S and D), complexity conditions (first- and second-order), and participant groups, computed for each electrode across the entire time window analyzed, using a 200 ms pre-S1 onset baseline. In red are marked the waveforms elicited by the electrodes selected for analysis (F3, F4, Fz, C3, C4, and Cz). Between bars appear the latencies selected to analyse the CNV and PINV components.

**Figure 3.**
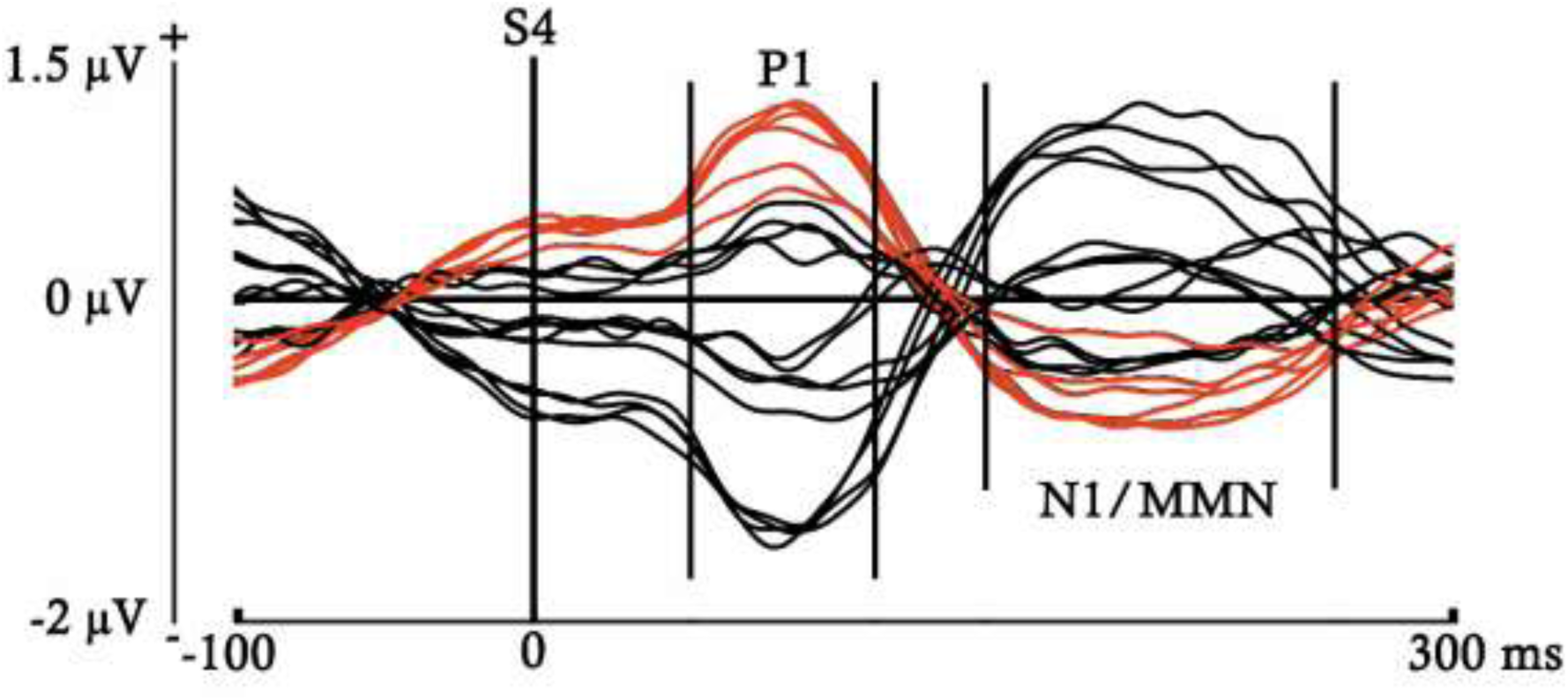
ERP averages obtained for both trial types (S and D), complexity conditions (first- and second-order), and participant groups, computed for each electrode from S4 onset to 300 ms, using a 100 ms pre-S4 onset baseline. In red are marked the waveforms elicited by the electrodes selected for analysis (F3, F4, Fz, C3, C4, and Cz). Between bars appear the latencies selected to analyze the P1 and MMN components. The N1/MMN label indicates that the MMN was embedded within a latency similar to that of the N1 component.

Figure 2 displays the entire interstimulus interval (0 to 2900ms post-S1), using a baseline of 200ms prior to S1, in order to identify components with longer temporal duration. In contrast, Figure 3 shows the voltage elicited during a shorter period following the onset of S4 (0 to 300ms post-S4), with a baseline of 100ms prior to S4, aiming to detect early components with shorter durations.

An artifact rejection voltage limit of ±130 mV was applied to the EEG in both the epochs selected to display Figures 2 (from 0 to 2900ms post-S1) and 3 (from 900 to 1200ms post-S1), and in all the epochs analyzed in this study (described below). This voltage rejection threshold is higher than that typically used in adults due to the higher spontaneous EEG amplitude observed in the pediatric population. This factor could lead to an excessively high number of rejected trials, potentially impairing the statistical power required to ensure optimal data quality. In this manner, all the epochs that exceeded ±130 mV in the time windows selected for each component, in any channel, complexity condition, and type of trial, was rejected (Table 2 and 3).

**Table 2.**
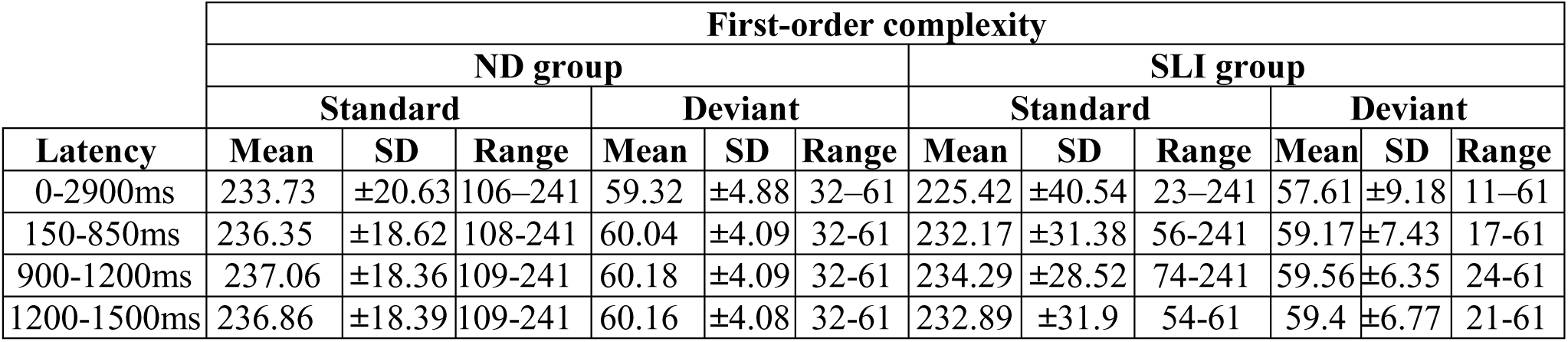
Accepted trials after voltage artifact rejection (first-order condition). The table presents the mean, standard deviation, and range of accepted trials applying an artifact rejection threshold of ±130 mV, for both participant groups and trial types, within the first-order complexity condition. Data are reported for four analyzed time windows: 0-2900ms (used for plotting the ERP grand-average); 150-850ms (selected for the CNV component); 900-1200ms (chosen for both the P1 and the MMN components); and 1200-1500ms (designated for the PINV component). The mean and standard deviation (SD) were measured in microvolts (µV).

**Table 3.**
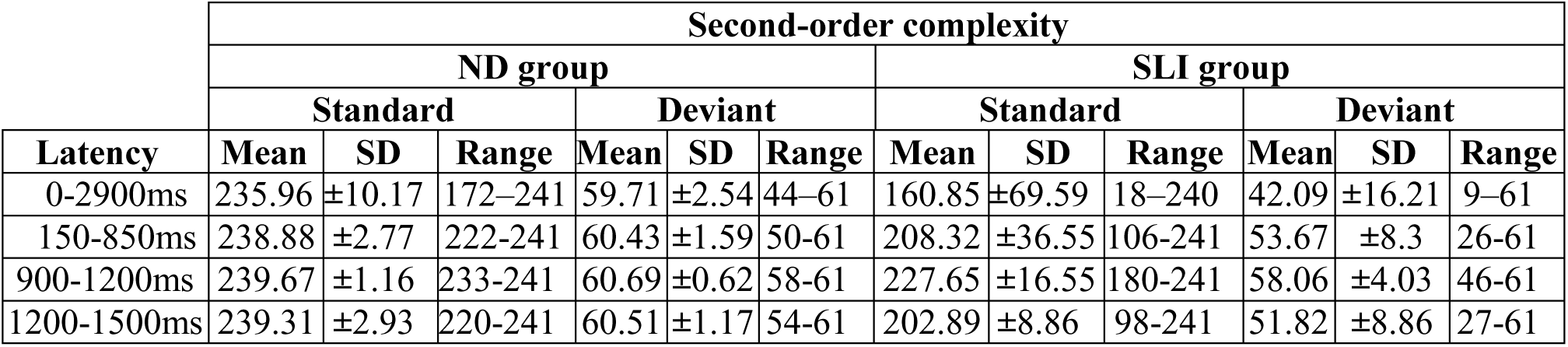
Accepted trials after voltage artifact rejection (second-order condition). The table presents the mean, standard deviation, and range of accepted trials applying an artifact rejection threshold of ±130 mV, for both participant groups and trial types, within the second-order complexity condition. Data are reported for four analyzed time windows: 0-2900ms (used for plotting the ERP grand-average); 150-850ms (selected for the CNV component); 900-1200ms (chosen for both the P1 and the MMN components); and 1200-1500ms (designated for the PINV component). The mean and standard deviation (SD) were measured in microvolts (µV).

The topographies of the mean ERPs for both participant groups and complexity conditions, for S and D trials, are plotted in Figure 4 for the time windows selected for each component analyzed (described above) to determine the electrodes most relevant for the analysis.

**Figure 4.**
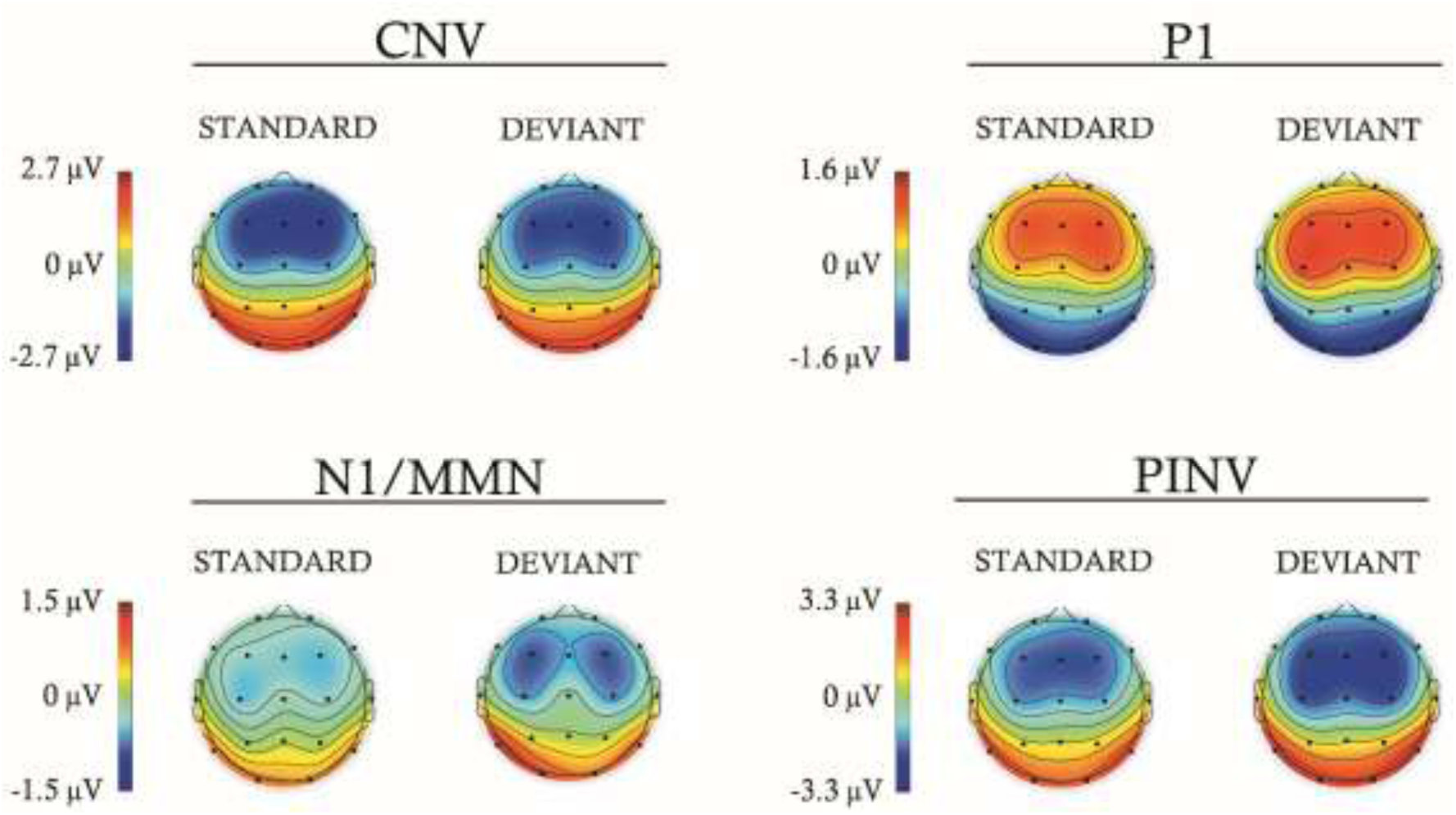
Topographies of the standard and deviant trials performed for the average of both complexity conditions (first- and second-order) and participant groups (ND and SLI), in each component analyzed (CNV, P1, N1/MMN, and PINV). The N1/MMN label indicates that the MMN was embedded within a latency similar to that of the N1 component.

The electrodes selected for analysis were F3, F4, Fz, C3, C4, and Cz, based on the higher amplitudes observed in Figure 4 (above) and the typical involvement of the frontocentral region in the components analyzed in this study (see above), as described in the previous literature (Sharma et al., 2005; Elbert et al., 1970; Gómez et al., 2019; Näätänen et al., 2012). A slow negativity from 150ms post-S1 to the S4 onset was selected for analysis (see Figure 2), with time window from 150 to 850ms post-S1 with a baseline of 200ms. This component would be interpreted as a CNV suggesting that it has been elicited by the expectancy generated by the S1 (considered as the warning signal) about the appearance of the S4 (considered the target) (Kononowicz et al., 2016; Arjona et al., 2017).

A progressive return to baseline was observed approximately from 350ms post-S4 to the end of the inter-stimulus interval (see Fig. 2). This component would be interpreted as a Post-Imperative Negative Variation (PINV), as it appears to reflect the ERP recovery of the negativity generated by the CNV (Wagner et al., 1996; Ruiz-Martínez et al., 2022). The time window was stablished from 345 to 600ms post-S4, because the effects was more clearly observed within this latency. A baseline of 200ms pre-S1 was used, due to the previously commented relationship of this component with the CNV.

An early and anterior positivity followed by a later and anterior negativity, possibly indexing P1 and N1/MMN component, respectively, can be observed in Figure 3 after the S4 onset. The time windows selected for analysis were from 50 to 110ms post-S4 in the case of the P1, and from 145 to 260ms post-S4 for the MMN. For both components the baseline was stablished 100ms previous to the S4. Note that in the figures displayed, the MMN latency was labelled as N1/MMN. The reason is that, although this time window was selected to analyze the MMN, presenting both trial types separately would correspond to the N1 component, theoretically, since the MMN is obtained by subtracting the S from the D. As a clarification for the results and discussion sections, when we refer to ERPs differences between S and D, we will use term MMN, whilst when referring to differences between first and second order complexities, which collapse S and D trials, we would use the term N1/MMN.

Four linear mixed-models analyses were computed on the mean voltage of the ERPs in the selected electrodes and time windows of the CNV, P1, MMN, and PINV components, considering the subjects as a random factor and the trial type (S or D), complexity (first- and second-order complexity), electrodes (F3, F4, Fz, C3, C4, and Cz), and participant group (ND and SLI) as fixed factors. Additionally, interactions between the participant groups and all other fixed factors were analyzed, which would be considered the more relevant to test our hypotheses. In the case that any interaction was significant, additional analyses were computed. These analyses compared the standard vs. deviant trials within each group, or the complexity paradigms between both subject groups, depending on whether the interaction with participants was significant for the trial type or the complexity paradigm, respectively. The gender and age in days of each participant were also included as fixed factors to assess and control their potential influence on the ERP components.

The ERPs for both type of trials and complexity conditions were displayed for both participant groups across the entire time window from the onset of the S1 to the end of the inter-stimulus interval, as well as for the 300ms post-S4, in order to better illustrate the effects described in the results.

## RESULTS

After a clearly formed auditory P1, MMN emerged as a modulation of the N1 component. Both components were observed in the ERP grand-average (Figures 5 and 6). Additionally, a CNV component was induced during the presentation of the four tones that composed the stimuli sequences (Figure 5). The negativity generated by the CNV is followed by a recovery to the baseline, starting about 350ms after the S4 onset, which was categorized as a PINV component.

**Figure 5.**
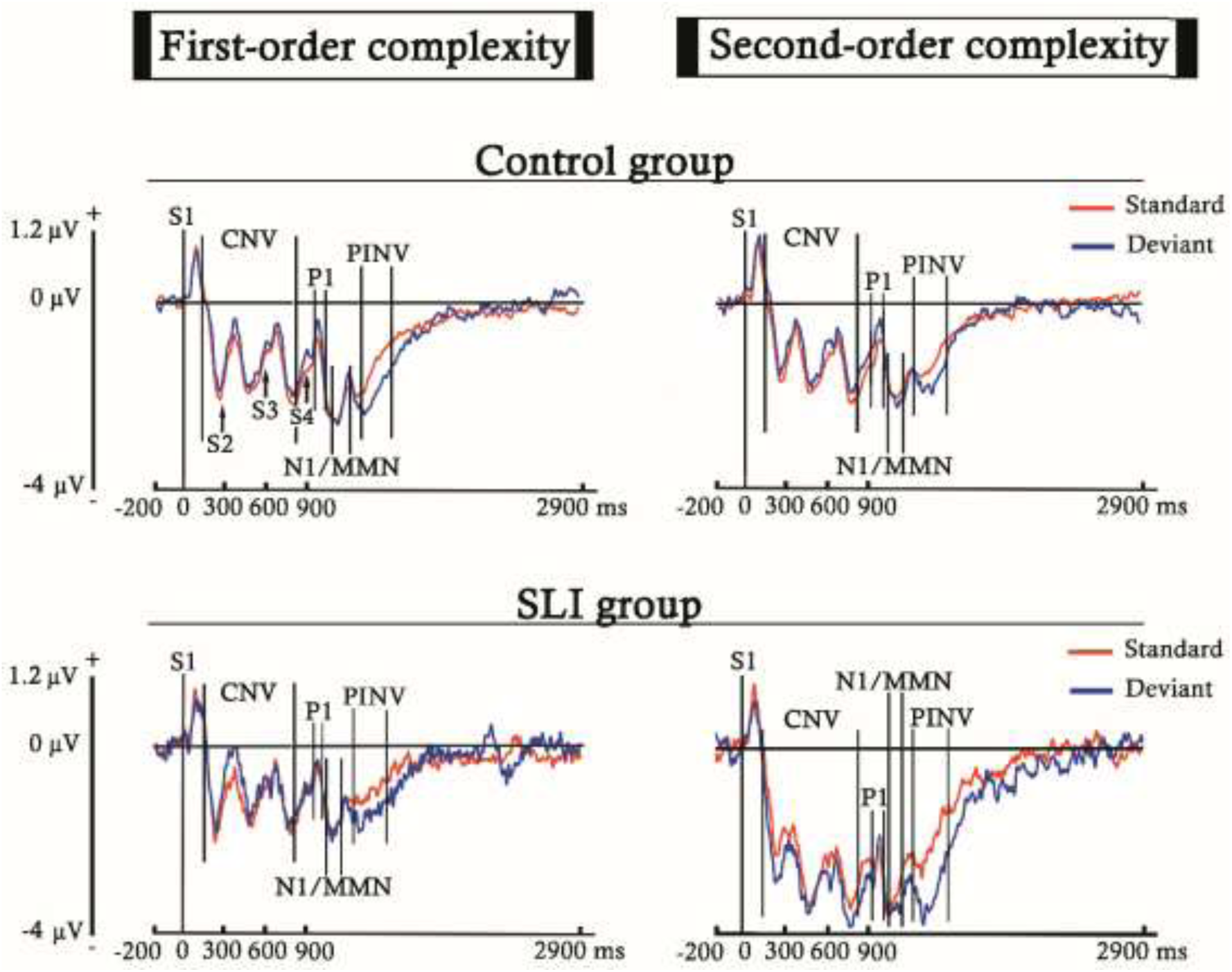
Event-related potentials (ERPs) elicited by the standard and deviant trials, for both the first- and second-order complexity paradigms, in the ND and SLI groups. ERPs were obtained by averaging the six electrodes selected for analysis (F3, F4, Fz, C3, C4, Cz). The baseline was established before the S1 (from -200 to 0 ms), and the plotted time window corresponds to the entire stimulus sequence and the inter-stimulus interval. The arrows pointing to the waveforms indicate the onset of every stimulus according to its presentation order (Sx). Between bars are delimited the latencies selected for the components analyzed (CNV, P1, N1/MMN, and PINV). The N1/MMN label indicates that the MMN was embedded within a latency similar to that of the N1 component.

**Figure 6.**
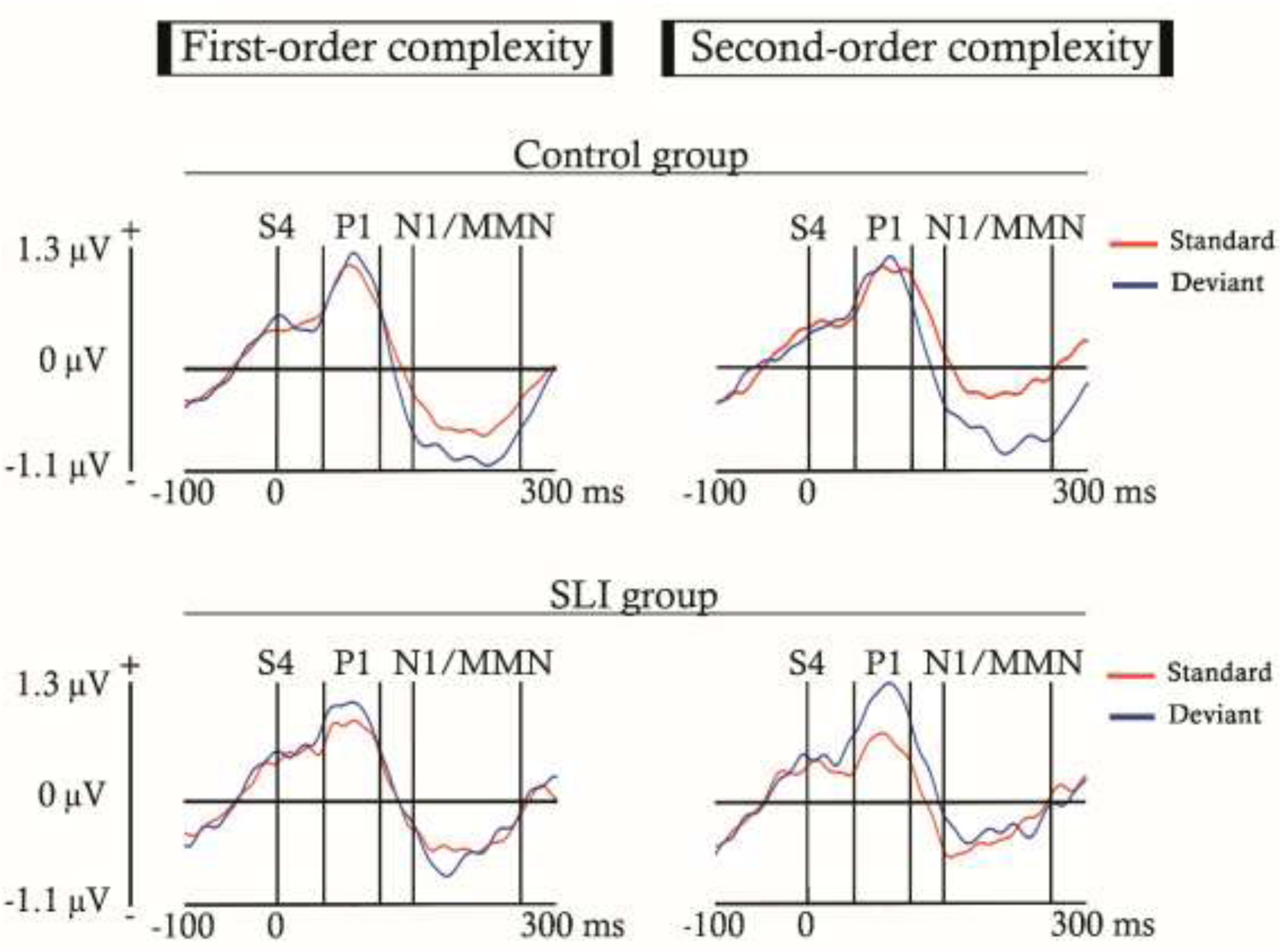
Event-related potentials (ERPs) elicited by the standard and deviant trials, for both the first- and second-order complexity paradigms, in the ND and SLI groups. ERPs were obtained by averaging the six electrodes selected for analysis (F3, F4, Fz, C3, C4, Cz). The baseline was established before the S4 (from -100 to 0 ms), and the plotted time window was 300 ms post-S4, allowing for clearer observation of the P1 and MMN components. Between bars are delimited the latencies selected for the components analyzed (P1 and N1/MMN). The N1/MMN label indicates that the MMN was embedded within a latency similar to that of the N1 component.

Below, we present the event-related potentials (ERPs) generated for each trial type, depending on the complexity condition and participant group. Figure 5 shows the ERP waveforms from the onset of the first stimulus of the sequence (S1) to the end of the inter-stimulus interval. In contrast, Figure 6 displays the same potentials, but restricted to the time window post-S4 selected for the P1 and MMN analyses, so that the effects described in these components can be more clearly observed.

The linear mixed model analysis (LMM) computed for the P1 component (Figure 6), with baseline located before S4, showed a significant effect for the interaction between the participant group and the type of trial (S or D) [F(1,4011) = 5.89, P = 0.015]. This difference between groups was due to the significant comparison between the S and D trial in the SLI group [F(1,1643) = 11.58, P = < 0.001], caused by the higher response elicited by the D (mean= 1.02 µV ± 2.31 SD) in comparison with the S (mean= 0.7 µV ± 1.68 SD). The increase in P1 amplitude was not significant in the ND group. Additionally, the electrodes analyzed (F3, F4, C3, C4, Cz, and Fz) also presented significant differences between them [F(1,4011) = 5.5, P = 0.019].

The analysis computed for the MMN, with baseline located before S4 (Figure 6), showed significant effects for the type of trial and its interaction with the participant groups, besides a significant result for the complexity paradigm condition (first vs. second-order).

The difference between the type of trial (S and D), is explained by a higher amplitude of the D trial (mean= -0.63 µV ± 2.37 SD) respecting the S (mean= -0.39 µV ± 1.7 SD). The differences between the first and second-order complexity is caused by a higher activity induced by the simplest auditory design (mean= -0.63 µV ± 2.02 SD), in comparison with the most complex (mean= -0.4 µV ± 2.1 SD). On the other hand, the interaction between the participant group and the type of trial is explained by a significant difference showed by the ND group [F(1,2396) = 31.6, P = < 0.001], in contrast to the SLI group. The difference between the type of trial found in the ND group was caused by a higher activation for the D (mean= -0.79 µV ± 2.21 SD) regarding the S (mean= -0.38 µV ± 1.56 SD). These results together suggest that while the D vs. S comparison in the SLI group is indexed by the P1 component, in the ND group it is indexed by the MMN.

The results obtained for the LMM computed for the PINV component showed significant effects for the complexity paradigm condition, as well as for its interaction with the participant group. Moreover, the group condition and the age measured in days also presented significant results.

The main effects observed for the complexity paradigm and the group were resulted from the higher amplitude generated by the second-order complexity (mean= -2 µV ± 3.13 SD) and the SLI group (mean= -1.93 µV ± 3.49 SD), in comparison with the first-order (mean= -1.56 µV ± 2.57 SD) and the ND group (mean= -1.68 µV ± 2.36 SD), respectively. The significant result obtained for the age measured in days in this component was caused by a progressively increasing negativity associated with increasing participant age (β = -0.08, t = 5.17, p <0.001).

The interaction between the complexity paradigms and the participant groups was due to a significant difference between the ND [F(1,84) = 4.25, P = 0.042] and SLI participants [F(1,83) = 14.6, P = <0.001] in both the first-order and second-order complexity conditions, although differences were in opposite sense in both groups. The ND group showed greater activation for the first-order complexity (mean= -1.79 µV ± 2.43 SD) respecting the SLI participants (mean= -1.56 µV ± 2.29 SD). In contrast, the SLI exhibited higher activation in the second-order condition (mean= -2.67 µV ± 3.99 SD), in comparison with the ND group (mean= -1.56 µV ± 2.29 SD). The increased activity observed in the clinical group for the more complex paradigm may reflect difficulties in processing stimuli, possibly accompanied by a sense of overload.

The analysis performed for the CNV component revealed significant effects for the participant groups, the type of trials and the age in days, as well as the complexity paradigm and its interaction with the participant groups.

The significant results obtained for the participant groups and the type of trial were caused by a higher activity elicited in the SLI group (mean= -1.79 µV ± 2.81 SD) and the S trial (mean= -1.58 µV ± 2.1 SD), compared to the ND group (mean= -1.31 µV ± 1.86 SD) and the D (mean= -1.44 µV ± 2.48 SD), respectively. The significant difference between both complexity paradigms was caused by a greater response generated by the second-order condition (mean= -1.84 µV ± 2.49 SD), in comparison with the first-order (mean= -1.17 µV ± 2.04 SD). Similar to the PINV component, the effect observed for the age (measured in days) in the CNV was due to an increase in negativity associated with increasing age (β = -0.06, t = -4.19, p <0.001).

On the other hand, the interaction with the participant groups was driven by a significant effect found in the second-order condition [F(1,83) = 19.94, P = < 0.001], but not in the first-order. This difference between the participant groups in the most complex paradigm was due to a greater activity recorded in the SLI group (mean= -2.62 µV ± 3.07 SD) compared to the ND group (mean= -1.33 µV ± 1.86 SD). The results obtained for this component would support the hypothesis that the second-order complexity condition induces cognitive overload in the SLI participants. This would also be reinforced by the greater overall activation shown by the SLI participants compared to the ND group.

As the difference in the PINV amplitude between the S and D trials was visually evident in both groups and complexity conditions (Figure 5), an *ad hoc* analysis with the electrodes and trial type as unique factors was conducted. The results indicated a significant effect for the electrodes recorded [F(1,4015.74) = 42.735, P = < 0.001], and also for the type of trial presented [F(1,4015.74) = 6.805, P = < 0.009] caused by higher PINV negativity in D (mean= -2.05 µV ± 3.25 SD) with respect to S trials (mean= -1.51 µV ± 2.41 SD). These results would suggest a still-developing predictive processing mechanism in this component, whose effects would be lost in the complete analysis as they are distributed across the other factors.

Finally, all the results presented were included in the following table to facilitate their visualization.

## DISCUSSION

The present study, which is grounded in the predictive coding theory (Friston, 2005; Winkler & Czigler, 2012), compared the psychophysiological responses of children diagnosed with Specific Language Impairment (SLI) and a ND group without neurological disorders while presenting two passive auditory paradigms of varying complexity. The aim was to explore whether a deficit or alteration in predictive neural mechanisms could underlie the language development difficulties observed in SLI, as suggested by the statistical learning model proposed for this disorder (Lammertink et al., 2014). The experimental paradigm was designed in order to stress the predictive mechanisms. Ultimately, the findings may support the development of a child-friendly protocol for aid in early diagnosis.

As proposed in the first study hypothesis, a stronger psychophysiological response was observed for the deviant (D) compared to the standard (S) trial, reflected in the MMN for the ND group (Table 4) and in the P1 component for the SLI participants (Fig. 6). This effect suggests that both groups exhibit predictive processing and then prediction error, regardless of complexity, as they respond more strongly to deviations from the extracted patterns. However, the ND group would show a more typical development of the neural mechanisms involved in prediction, which the MMN index engaged in prediction error processing would support, as observed in healthy adults and children using the same (Ruiz-Martínez et al., 2022) or similar experimental designs (Tervaniemi et al., 1994; Muñoz-Caracuel et al., 2024).

**Table 4.**
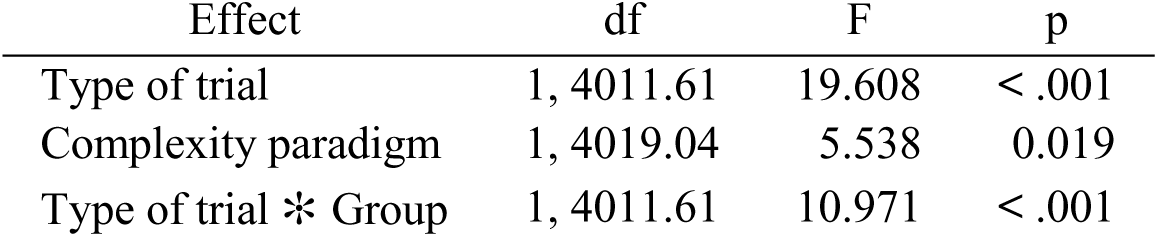
Significant LMM results for MMN. The table presents the significant results obtained from the Linear Mixed Model (LMM) analysis of the Mismatch Negativity (MMN) component.

The MMN is more associated with prediction error and deeper cognitive and attentional mechanisms involved in extracting predictive patterns (Winkler, 2007; Winkler & Czigler, 2012; Paavilainen et al., 2013), which has been related to the correct language development (Cheour et al., 2000; Wang et al, 2012; Paquette et al., 2013), or its impairment when the component is reduced or absent (Shafer et al., 2005; Datta et al., 2010; Kujala et al., 2017). For this reason, the fact that the SLI participants showed a discrimination effect between S and D trials in the P1 component instead of in the MMN would suggest a prediction processing, based on more basic sensory encoded features of auditory stimulus (Herrmann & Knight., 2001; Pratt, 2011). This role of the P1 component in deviance detection is aligned with the idea of a hierarchically organized novelty and deviance detection system in the human auditory system, which would present early indices of deviance detection preceding the MMN, which would be more involved in the extraction of abstract regularities (Slabu et al., 2010, Grimm et al., 2011; Grimm & Escera, 2012).

Additionally, the fact that prediction process in SLI was observed in the P1 component, rather than the MMN (Fig. 6), may reflect an impairment in the typical development of the brain regions involved in predictive coding. The typical maturation of the central auditory system shows a progressive P1 reduction from the childhood (Ponton et al., 2002; Mahajan & McArthur, 2012; Sharma et al., 2005; Wunderlich et al., 2006), while the MMN increase to the adulthood, in a process in which the neural generators associated with the MMN gradually acquire greater functional relevance. (Gomot et al., 2000; Kushnerenko et al., 2002; Guzzetta et al., 2011). Although both components originate in the primary auditory cortex (specifically in Heschl’s Gyrus) (Fitzroy y col., 2015; Ruhnau y col., 2011), the MMN also involves frontal generators related to attention-switching or orienting that is triggered by the change detection mechanism (Giard et al., 1990; Rinne et al., 2000; Garrido et al., 2009). In contrast, the frontal activity observed for the P1 (see Fig. 4) would result from thalamocortical projections (Ponton & Eggermont, 2001). Thus, the involvement of P1 in predictive processing in children with SLI, instead of the MMN, may reflect a psychophysiological compensation mechanism for an impaired or atypically slow maturation of the frontal MMN generators.

The higher amplitude observed for the S trial in the CNV component would suggest that both groups correctly allocated attention and anticipated upcoming stimuli. This effect is consistent with previous findings, showing that CNV amplitude increases as the predictability of a stimulus rises (Barry et al., 2007; Arjona & Gómez, 2014; Duma et al., 2020). In our study, this effect is likely due to the task design, in which the probability of an S trial is higher after a D trial or complete after two consecutive D-D trials, enhancing the anticipatory processing. On the other hand, the PINV did not show a significant difference between both type of trials until, motivated by the effect visually observable in Figure 5, an additional *ad hoc* LMM was computed, including only the electrode and trial type factors. This analysis revealed a significant effect for both factors, caused by a higher response elicited by the D, primarily in frontal areas (Figure 4). These results would suggest the involvement of a still-developing prediction mechanism for this component, as the effects were not sufficiently robust to withstand the influence of other factors.

Regarding the second hypothesis of the study, the N1/MMN showed a higher amplitude for the first-order paradigm when compared to the second-order condition (Table 4), whereas the P1 component did not show statistically significant differences between the first and second-order paradigms (Fig. 6). In line with this, the MMN sensitivity to paradigm complexity has been more frequently reported in the literature (Saarinen et al., 1992; Paavilainen et al., 2013; Paavilainen et al., 2018). The higher response to the simplest paradigm in both groups contrasts with the effects described in adults using the same experimental design, in which participants showed a greater activation for the second-order condition (Ruiz-Martínez et al., 2022). This would suggest that, although the neural mechanisms involved in extracting predictions from varying complexity patterns may differ between children and adults, SLI individuals do not show altered complexity processing in the N1/MMN.

The higher response for the first-order condition could reflect that the frontal N1/MMN generators, involved in analyzing changes in complex patterns (Korzyukov et al., 2003; Schröger et al., 2007), are still in a maturation process, given the typically slow development of the frontal areas (Bunge et al., 2002; Ivry & Knight, 2002). This effect would be observed as a reduced activity in the most complex paradigm, in which N1/MMN generators should be in higher demand. This result is in line with the findings described by Gumenyuk et al. (2003), who also pointed to the slow maturation of the frontal lobes to explain the lower MMN response elicited in frontal areas, in children aged 8 to 14, by the hard condition of a study consisting of two auditory patterns of different complexity. Secondarily, the increased amplitude observed in the first-order complexity condition supports the predictive coding over the adaptation theory. So that, under the adaptation account, the reduced number of frequencies should lead to lower activation, due to repeated exposure to the same tones which would cause neural habituation (May & Tiitinen, 2009; O’Reilly & O’Reilly, 2021).

Addressing the third proposed hypothesis, the effects observed for the comparison between both complexity conditions in the PINV also revealed significant effects in opposite directions between groups (Table 5; Fig. 5). So that while ND subjects showed a higher response for the first-order paradigm, the SLI participant presented a greater amplitude for the second-order condition. The absence of differences between the type of trial in this component, when all the factors analyzed were included, and the increased negativity associated with increasing age could imply an immature development of the update-predictive functions for the PINV in this age range (from 3 to 11 years old). This proposal would be supported by the findings described in adults applying the same or similar experimental design, in which they showed significant effects both in the comparison between the S and D trials (Muñoz-Caracuel et al., 2024), as in the first- and second-order conditions (Ruiz-Martínez et al., 2022).

**Table 5.**
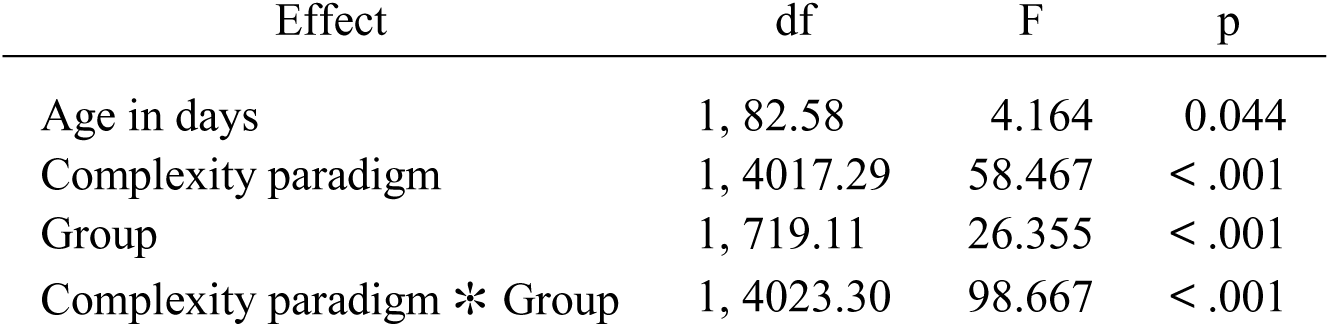
Significant LMM results for PINV. The table presents the significant results obtained from the Linear Mixed Model (LMM) analysis of the Post-Imperative Negative Variation (PINV) component.

For these reasons, the PINV observed in the ND group could merely reflect the progressive recovery of the CNV, maintaining the amplitude difference between both complexity conditions previously triggered by the MMN. Meanwhile, in the SLI group, the higher response to the second-order complexity, along with the absence of adequate predictive error, as indexed by P1 but not by MMN, could reflect cognitive overload. This presumable overload would be caused by the greater abstraction demands of the paradigm, which leads to difficulties integrating responses into a symbolic framework and/or a perceived lack of control over an aversive (in this case, excessive) stimulation, as described for this component (Delaunoy et al., 1978; Kathmann et al., 1990; Diener et al., 2009).

A similar effect, during the second-order complexity paradigm, was observed for the CNV component in SLI participants (Table 6; Fig. 5), which possibly would indicate a greater anticipatory effort (Kropp et al., 2001; Kononowicz & Penney, 2016; Fattapposta et al., 2024). This result, along with the overall increased activity recorded in this group (similar to the effects described in adults by Plante et al. in 2017), supports the presence of both cognitive overload and enhanced effort to reassess stimulus patterns in children with SLI in response to predictive coding paradigms.

**Table 6.**
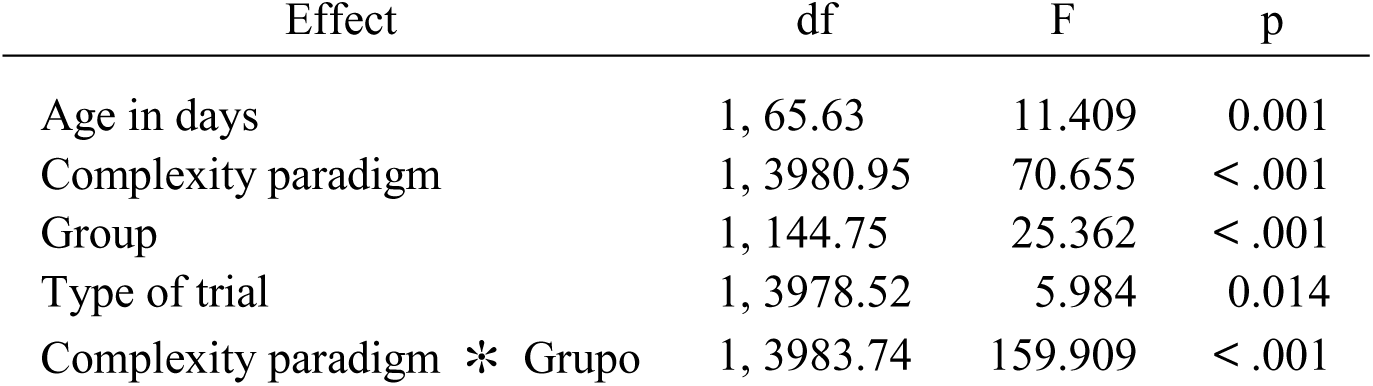
Significant LMM results for CNV. The table presents the significant results obtained from the Linear Mixed Model (LMM) analysis of the Contingent Negative Variation (CNV) component.

## CONCLUSIONS

The current study evaluated the statistical learning model explanation for language difficulties in children with SLI, using for this goal a two passive auditory paradigms of varying complexity framed within the predictive coding theory. The SLI participants showed an absent MMN and a higher P1 response to D tones, which may indicate immature or impaired development of frontal MMN generators and, possibly, a compensatory mechanism involving earlier auditory processing areas. On the other hand, the increased PINV and CNV amplitudes observed in the SLI participants during the most complex condition could suggest cognitive overload and heightened effort to reassess stimulus patterns in this group.

Additionally, both groups responded more strongly to the first-order paradigm compared with the second-order condition, which could suggest that the frontal regions implicated in abstract pattern extraction are still maturing. Finally, all these findings (see Table 7) would support the statistical learning model as a valid approach for researching the neural basis of the SLI disorder. This could promote the foundation of an early detection protocol through EEG, given its non-invasive, child-friendly, and passive features.

**Table 7.**
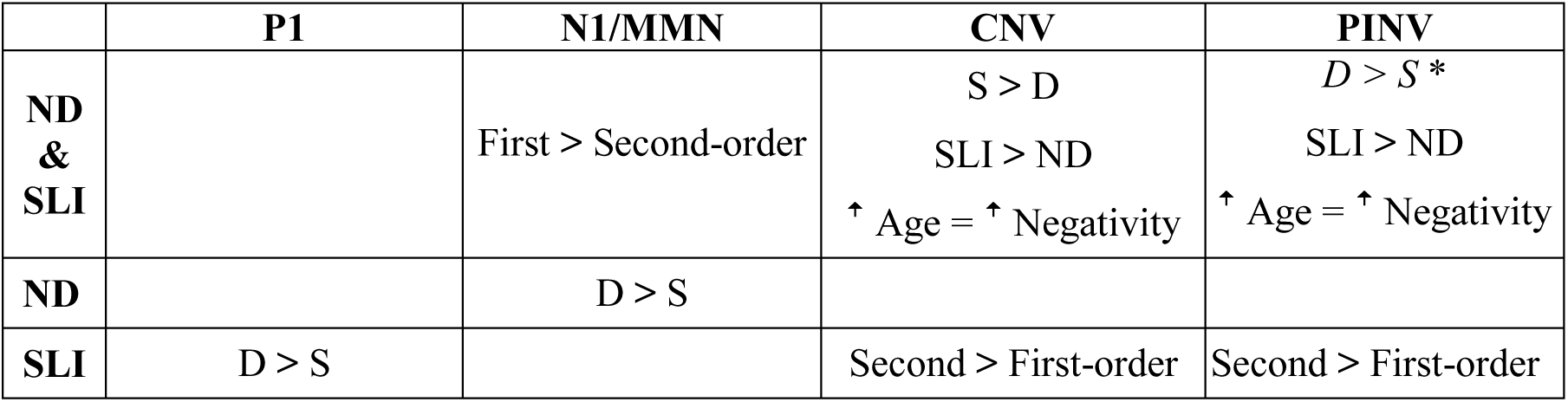
Summary of key results by component and group. Summary of all results of interest obtained in the study for each component analyzed. The row ND & SLI shows the results observed in both groups, whereas the rows ND and SLI present the results obtained for each group separately, specifically after an interaction between a factor and the group. The upper arrow indicate that negativity increased with age. *This result was obtained when computing the LMM only with the factors electrodes and trial type.

## LIMITATIONS

Between the limitations of present report, the variety of diagnostic measures preclude the possibility of establishing correlational relationships between EEG and language performance.

## DATA AVAILABILITY STATEMENT

Data and code related with the present study is available under reasonable request to the corresponding author

## ACKNOWLEDGMENTS

We are grateful to the children who took part in the study and their families for their trust and collaboration throughout the process. We also extend our appreciation to the Unidad de Desarrollo Infantil y Atención Temprana (UDIATE), associated with the Victoria Eugenia Hospital of the Spanish Red Cross, for their important support in referring children for enrollment. The authors have carefully reviewed and revised the manuscript and accept full responsibility for its content.

